# Computational model of the full-length TSH receptor

**DOI:** 10.1101/2022.07.13.499955

**Authors:** Mihaly Mezei, Rauf Latif, Terry F Davies

## Abstract

The receptor for thyroid stimulating hormone (TSHR), a GPCR, is of particular interest as the primary antigen in autoimmune hyperthyroidism (Graves’ disease) caused by stimulating TSHR antibodies. To date, only one domain of the extracellular region of the TSHR has been crystallized. We have now generated a model of the entire TSHR by merging the extracellular region of the receptor, obtained using artificial intelligence, with our recent homology model of the transmembrane domain, embedded it in a lipid membrane solvated it with water and counterions, and performed 1000ns Molecular Dynamic simulations on it.

The simulations showed that the structure of the transmembrane and leucine-rich domains were remarkably constant while the linking region (LR), known more commonly as the “hinge region”, showed significant flexibility, forming several transient secondary structural elements. Furthermore, the relative orientation of the leucine-rich domain with the rest of the receptor was also seen to be variable. These data suggest that this linker region is an intrinsically disordered protein (IDP). Furthermore, preliminary data simulating the full TSHR model complexed with its ligand (TSH) showed that (a) there is a strong affinity between the linker region and TSH ligand and (b) the association of the linker region and the TSH ligand reduces the structural fluctuations in the linker region.

This full-length model illustrates the importance of the linker region in responding to ligand binding and lays the foundation for studies of pathologic TSHR autoantibodies complexed with the TSHR to give further insight into their interaction with the flexible linker region.

## Introduction

The thyroid-stimulating hormone receptor (TSHR) on the surface of thyrocytes is an important regulator of thyroid growth, development, hormone synthesis and secretion. It is also the primary target of autoantibodies in Graves’ disease (autoimmune hyperthyroidism) (1). From cloning, sequence analysis, partial crystallization and biochemical studies this GPCR has been deduced to be made of a large ectodomain (ECD) and membrane-bound signal transducing transmembrane domain (TMD) (2,3). The ECD is further divided into a leucine rich domain (LRD) forming a curved structure which is linked to the TMD by a 130 amino acid (AA) linker region (LR) known commonly as the “hinge region” (AA280-410). Unique to the TSHR is a large 50 amino acid cleavage region (AA316-366) located within the LR that is proteolytically degraded leaving a cleaved ectodomain thought to be tethered to the TMD via 3 cysteine bonds (4,5) and sometimes referred to as the C peptide.

Crystallization studies (6,7), besides producing crystal structures for the LRD, have shown that antibodies, which either stimulate or block TSHR signaling, bind to the LRD when the receptor is conformationally correct and can compete for TSH binding. In contrast, “neutral” antibodies to the TSHR which do not initiate a traditional signal (8), nor inhibit TSH binding, predominantly, but not exclusively, bind to linear epitopes in the LR (9). Although the partial LRD structure has been determined with x-ray crystallography (10) no experimental structure has been found for the LR and, until recently, only partial models have been proposed (11–13). On the basis of the immune response to the TSHR we, and others, have suggested that the LR is not an inert scaffold but rather an important ligand-specific structural and functional entity (14) but its structure has not been examined in the context of the full-length receptor. However, the recent success of the AI-based Alphafold2 (15) program lead us to believe that it might be possible to generate a full-length receptor structure by combining the LRD-LR structure generated by Alphafold2 (that includes a structural model of the LR region but no TMD and for which neither experimental nor homology models are available) with our recently published model of the TSHR TMD (named TRIO) (16). This full-length model could then be enhanced and verified with molecular dynamics simulation (MD). We can now report a successful computer-based approach to obtain insight into the LR allowing us to complete a full-length model of the TSHR. We have examined the behavior of this TSHR in a lipid-embedded, electro-neutral, aqueous environment by molecular dynamic simulation studies and showed that the LR is indeed an intrinsically disordered protein but can be stabilized by TSH ligand binding to the LRD.

## MATERIALS AND METHODS

### AI model of the LRD and LR

The coordinates of the structure of the LRD and LR domains of the human TSHR, residues 24 – 408, were downloaded from the Swissprot database (17); Uniprot #: P16473 and Swissprot file /P1/64/73. The first 23 residues, of which 21 residues formed the signal peptide, were not included in the model.

### Model of the TMD

Our previous work (16) detailed the MD trajectory of the TMD, residues 408 – 717 of the human TSHR. into three clusters using k-medoid clustering (18), performed by the program Simulaid (19). For the present work, we chose the representative structure of the largest cluster, forming during the second half of the MD trajectory. As before, the initial positions of internal waters were determined using grand-canonical ensemble Monte Carlo simulation (20), followed by circular variance (21) filtering and derivation of generic sites (22). The Monte Carlo simulation, as well as the circular variance and generic site calculations, were performed with the program MMC (23).

### Formation of a full-length TSH model by combination of the Alphafold2 LRD and LR model with the TRIO TMD model

The alphafold2 model of the TSHR ectodomain (LRD-LR) and the TRIO model have only one common residue – cysteine 408. First, the LRD-LR model was translated so that the Cα of the LRD-LR cysteine is at the position of the TRIO cysteine’s Cα position. As both models were already oriented along the membrane normal (the Z axis), the only degree of freedom left was rotation around the Z axis. In the next step a scan by 45° steps selected the angle region that minimized the volume of the enclosing rectangle, followed by generating conformations in 5° steps and obtaining the list of contact distances between the LR and the TMD (pairs of atoms are defined to be in contact if they are mutually proximal). Examination of the contacts narrowed down the likely conformation. The final choice was made after having examined visually (using the program VMD (24)) the form that resulted in the broadest contacts between the LR and the TMD. While this last step is admittedly an inexact operation, it is made with the understanding that small errors would be corrected during the MD equilibration. The coordinates of this initial model is available from the Dryad server at the URL https://doi.org/10.5061/dryad.rjdfn2zdp.

### Immersion in bilayer

The Charmm-Gui server (25) was used to immerse the full model of TSHR, including the internal waters, into a bilayer of DPPC molecules. The server also added a water layer as well as counterions (K^+^ and Cl^-^ ions), both to ensure electroneutrality and an ionic strength of 0.15 M to best represent physiological conditions. Periodic boundary conditions were applied using a hexagonal prism simulation cell. The system thus generated included inputs for a six-step equilibration protocol and inputs for the production run, all using the program NAMD (26).

### Molecular dynamics simulation

The simulations used the default parameters set by Charmm-gui. For the protein and the ions the pairwise additive Charmm36m force field (27) was used and water was represented by the TIP3P water model(28). Long-range electrostatics was treated with the Ewald method and VdW interactions used a cutoff of 12 Â, smoothly cut to zero starting at 10 Â. The simulations used 2 fs time steps and were run in the (T,P,N) ensemble.

### TSHR-TSH and TSHR-antibody complexes

The TSHR-TSH complex used for calculating clashes with the Alphafold2 model and for the start of the MD simulation was obtained based on the crystal structure of the Follicle-Stimulating Hormone (FSH) in complex of the LRD of FSH (PDB id 4ay9). In the first step the several TSH beta chain coordinates were generated based on the FSH beta chain coordinates in the FSHR-FSH complex using the program Modeller (29). Next, the model with that had the fewest clashes with the LRD was selected and used to replace the FSH beta coordinates, followed by aligning the LRD of the FSHR-TSH beta complex to the LRD of our full TSHR model. Finally, the beta chain of one structure from an earlier unpublished model of the TSH-LRD complex was aligned to the beta chain of the newly generated complex to add the TSH alpha chain to the model.

The TSHR-antibody complexes for the calculation of clashes were obtained by superimposing the LRD domains in the crystal structures of stimulating and blocking antibodies (PDB ids 3g84 and 2xwt) to the LRD domain of the Alphafold2 structure.

### Analyses

Most analyses were performed on the trajectories with the program Simulaid (19). Hydrogen bonds are defined by Simulaid as X···H-Y where X and Y are polar heavy atoms, the X···H-Y angle is above 120° and the X-Y distance is below threshold. Note that this definition ignores the actual charges thus it includes salt bridges as well; as is the case for several of the hydrogen bonds thus defined. The adequacy of the run length was verified by the saturation of the hydrogen-bond tracks. In other words, after a while the system did not form new hydrogen bonds, it only broke and reformed the existing ones. The variation of the shape of the LR was tracked by calculating the radius of gyration. The change in the relative orientation of the LRD with the rest of the protein was characterized by the angle between the first principal axis of the LRD and the Z axis and also tracked by observing the animated trajectory. Formation and unraveling of secondary structure elements in the LR was tracked with the DSSP algorithm (30).

## Results

### Initial full-length TSHR model and its clashes

**Figure 1** shows the structure of the full-length receptor that was obtained by combining the LRD-LR structure generated by the Alphafold2 AI system with the TMD structure from the TRIO model as detailed in Methods section. The LR is also shown in the enlargement, with the centers of five linear epitopes, which are immunogenic in a full length TSHR immunized mouse model (9), shown as spheres. Note that while the LR region has unique linear epitopes for TSHR antibodies of the “neutral” variety, the LRD is not devoid of such binding sites either, with at least 3 such linear epitopes (not represented here) seen in immunized mice (9).

**Figure 1:**
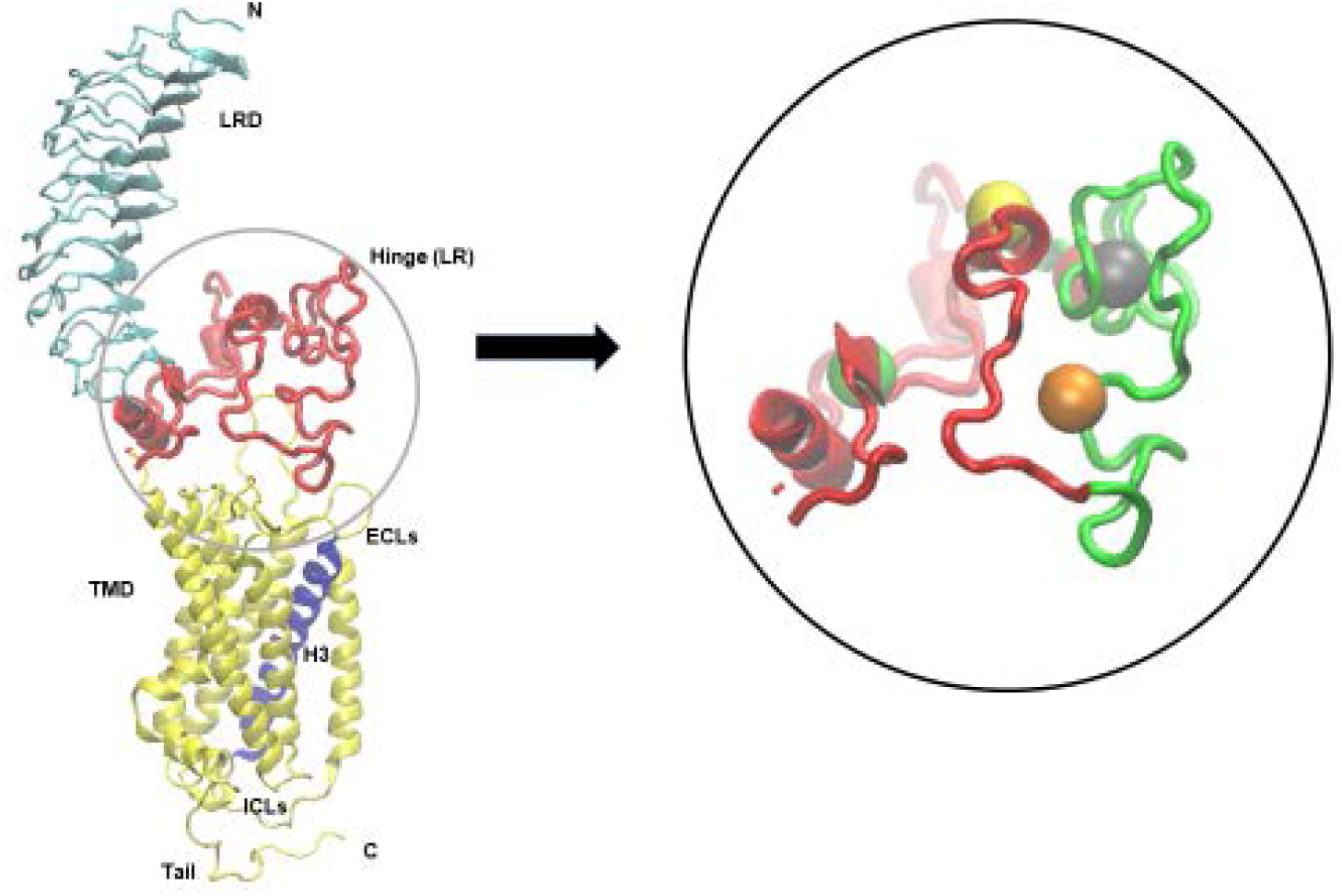
The initial model of the full-length TSHR (LRD: blue, LR: red, TMD: yellow) derived from combination of the LRD and hinge region of the Alphafold2 program and the TMD of our earlier “TRIO” model (16). Helix 3 of the TMD is shown in purple. The enlargement shows the LR with the unique 50 AA insert in green. In addition, this diagram shows the Cα atoms at the centers of our described LR epitopes (13) as spheres; epitope 313-324: red (partly obscured), epitope 322-342: gray, epitope 349-356: orange, epitope 377-391: yellow, and epitope 393-400: green.

However, this assembled full length TSHR structure showed significant LR clashes with TSH ligand and with TSHR antibodies (**Figures 2A-C**). A clash was here defined as a heavy atom distance less than 2.10 Â, 1.68 Â, and 1.65 Â, for atom pairs involving S, N or O, and C, resp. In particular, the number of LR heavy atoms clashing with TSH, blocking antibody (PDB id 2xwt), and stimulating antibody (PDB id 3g04) were 40, 68, and 101, resp. and involved 7, 11, and 18 residues, resp. Clearly, this approach showed marked hindrance of ligand and autoantibody binding so the model required further development.

**Figure 2:**
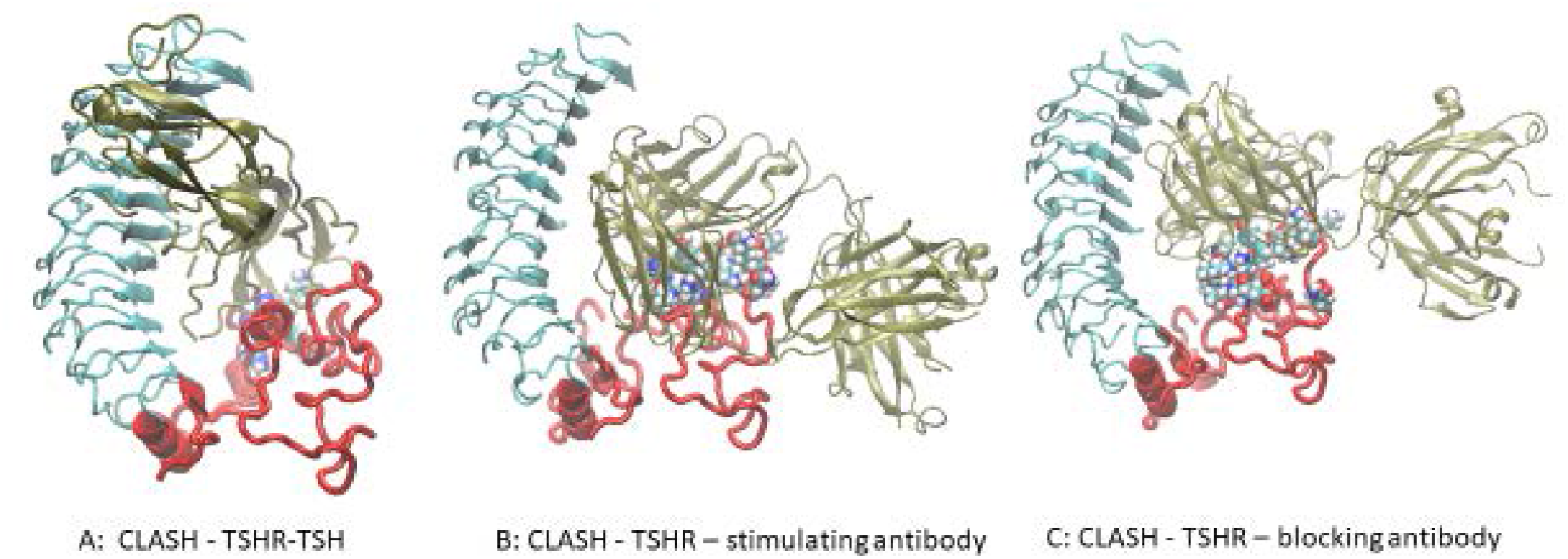
(A) The extracellular part of the full-length model from Figure 1 is shown in combination with the TSH ligand. The LR backbone is shown in red and several LR residues clashing with the TSH are shown as spheres (partly obscured). For clarity the TMD has been removed in this and subsequent illustrations. (B) Similarly, the LR model is shown clashing with a stimulating TSHR monoclonal antibody (MS-1) based on the crystal structure (PDB id 3g04) with even more clashes than with TSH. (C) Here the LR is clashing with a blocking TSHR monoclonal antibody (KI-70) based on the crystal structure (PDB id 2xwt) which once again shows many clashes.

### Inserting the TSHR model into a cell membrane

The combined model, including Monte Carlo-generated internal waters, was then sent to the Charmm-Gui server to be embedded in a DPCC lipid bilayer and immersed in water with counterions. The membrane-inserted, fully hydrated and neutralized system consisted of 177 and 176 DPPC molecules in the upper and lower layer, respectively, 148 K^+^ and 149 Cl^-^ ions and 54,412 water molecules, a total of 220,799 atoms in the simulation cell. The length of the periodic cell hexagon was 185.0 Â and the edge of the base hexagon was 66.4 Â. **Figure 3** shows the full simulation cell.

**Figure 3:**
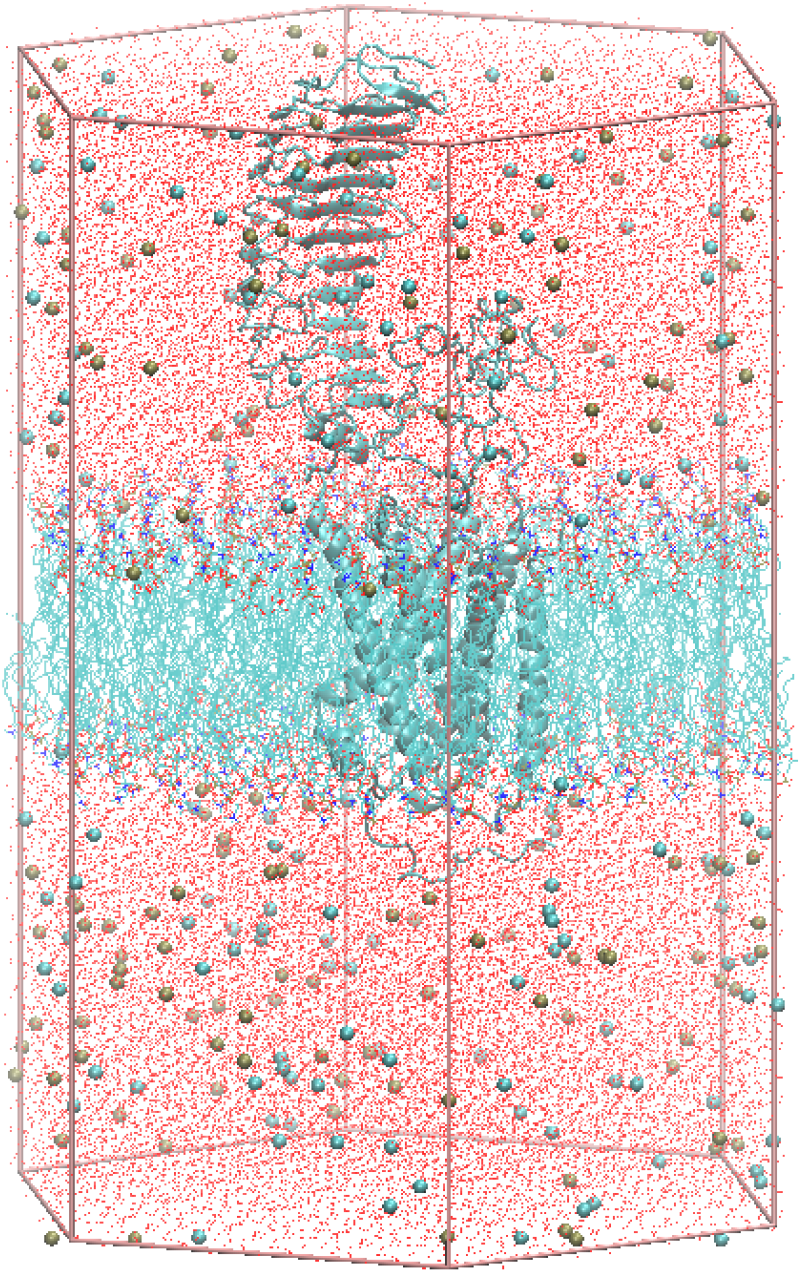
The full simulation cell prepared by Charmm-gui. The TSHR is shown in gray cartoon representation, lipids are shown as lines without hydrogens, ions as tan or cyan spheres representing K^+^ or Cl^-^ ions, resp., and the water oxygens as red dots. The hexagonal prism edges defined the initial simulation cell.

As the structures of the LRD have been experimentally determined by crystallography and we have described the TSHR-TMD in detail earlier (16), the analyses in this report are focused on extracting a potential structure of the entire LR from the MD simulation trajectory of the Alphafold2-based membrane embedded structure.

Animation (with the VMD software) of the simulation trajectory showed that (a) the LRD and TMD structures in the simulated complex did not show significant deviation from the earlier reports; (b) the LR structure generated using the Alphafold2 program had few secondary structural elements and showed significant fluctuation; and (c) the relative orientation of the LRD with the rest of the protein also fluctuated significantly during the simulation. This, therefore, offered the opportunity for finding conformations where the possibility of ligand and antibody binding did not clash with the LR. **Figure 4A** shows the 2D RMSD map of the LR over 2000 evenly spaced conformations. The RMSDs are calculated for the LR backbones. K-medoid clustering was performed asking for three clusters (as suggested by the 2D RMSD map) and the cluster representatives (the structure with the lowest maximum RMSD with the rest of the cluster members) were also extracted. These three representative structures of the LR are shown in **Figure 5** with the LR backbones of the three clusters in red and illustrating the unique 50 AA cleaved region.in green along with their simulation times.

**Figure 4:**
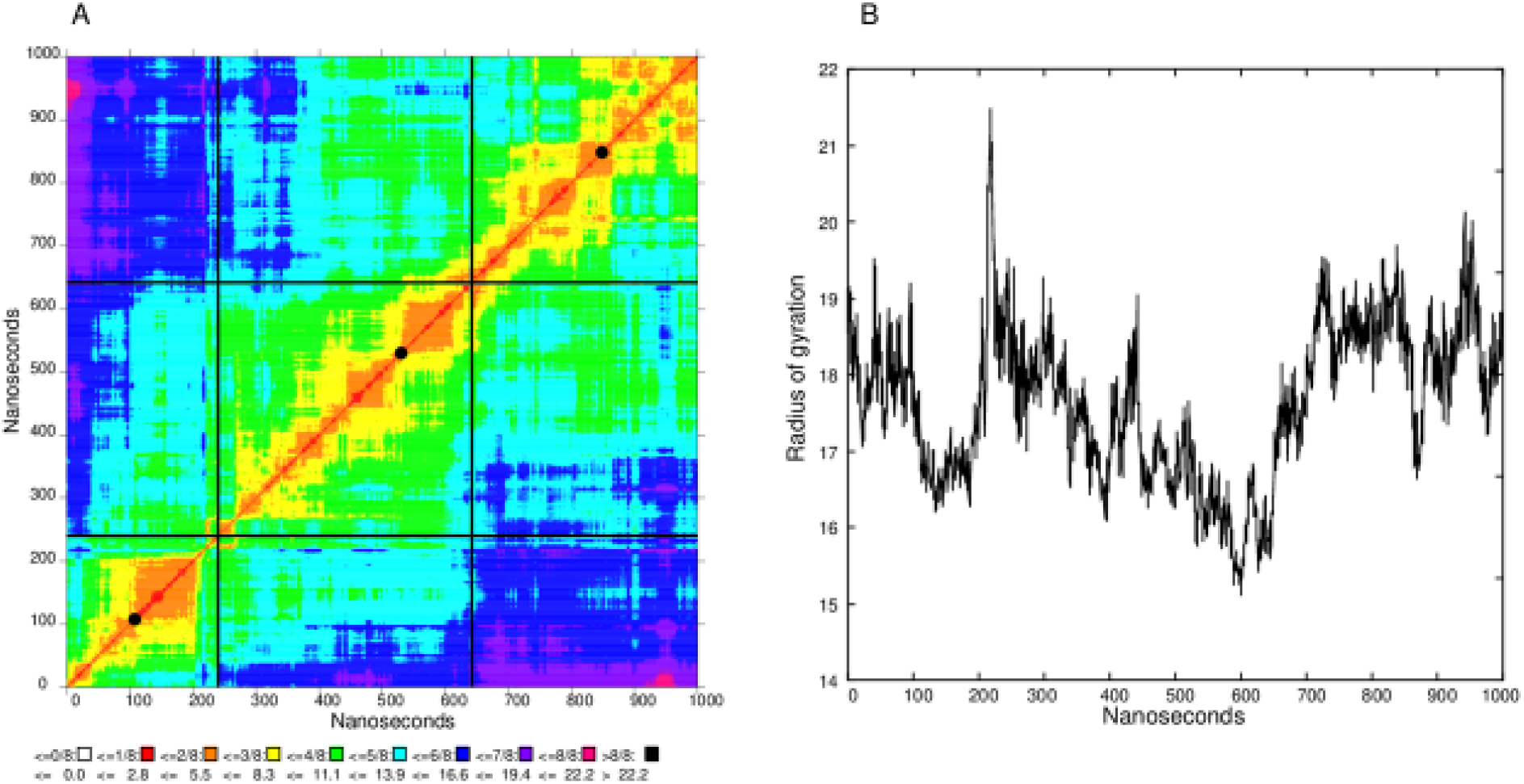
(A) 2D RMSD map of the LR during the 1000 ns simulation. Black lines delineate the three clusters and the black discs on the diagonal indicate the most representative structure. (B) The radius of gyration of the LR during the simulation.

**Figure 5:**
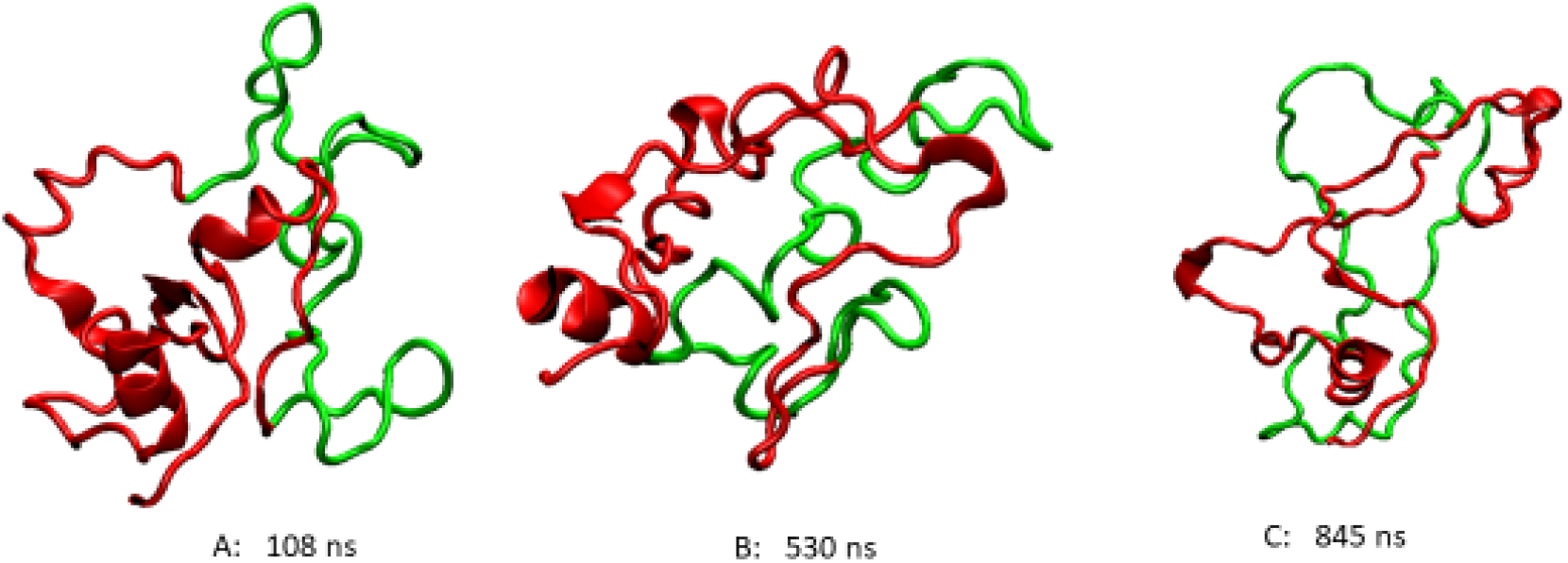
These three clusters are representative of the highly flexible structures of the LR backbone at different times during the 1000 ns simulation. The 50 AA cleaved segment is shown in green, the rest of the LR is in red.

### Instability of the LR

Examination of these backbones clearly showed that the LR does not form a well-defined stable tertiary structure. The radius of gyration *R*_g_, a measure of compactness, of the LR is shown in **Figure 4B** as a function of simulation time. It shows remarkable fluctuations with the range (the difference between the highest and lowest value) of *R*_g_ values being 6.4 Å. In contrast, the range of *R*_g_ values was only 1.6 Å for the larger LRD (not shown). The secondary structure of the LR was also tracked by the DSSP algorithm.

**Figure 6A** shows the secondary structural elements (SSE) found as the simulation progressed. Most SSEs are helices but, remarkably, in the 700-900 ns range several beta sheets formed and then dissolved. While a helix at the N terminal (residues 280-290) persisted throughout the calculations, **Figure 6A** also shows that all the other transient helices were seen to unwind or form only in the later stages of the simulation.

**Figure 6:**
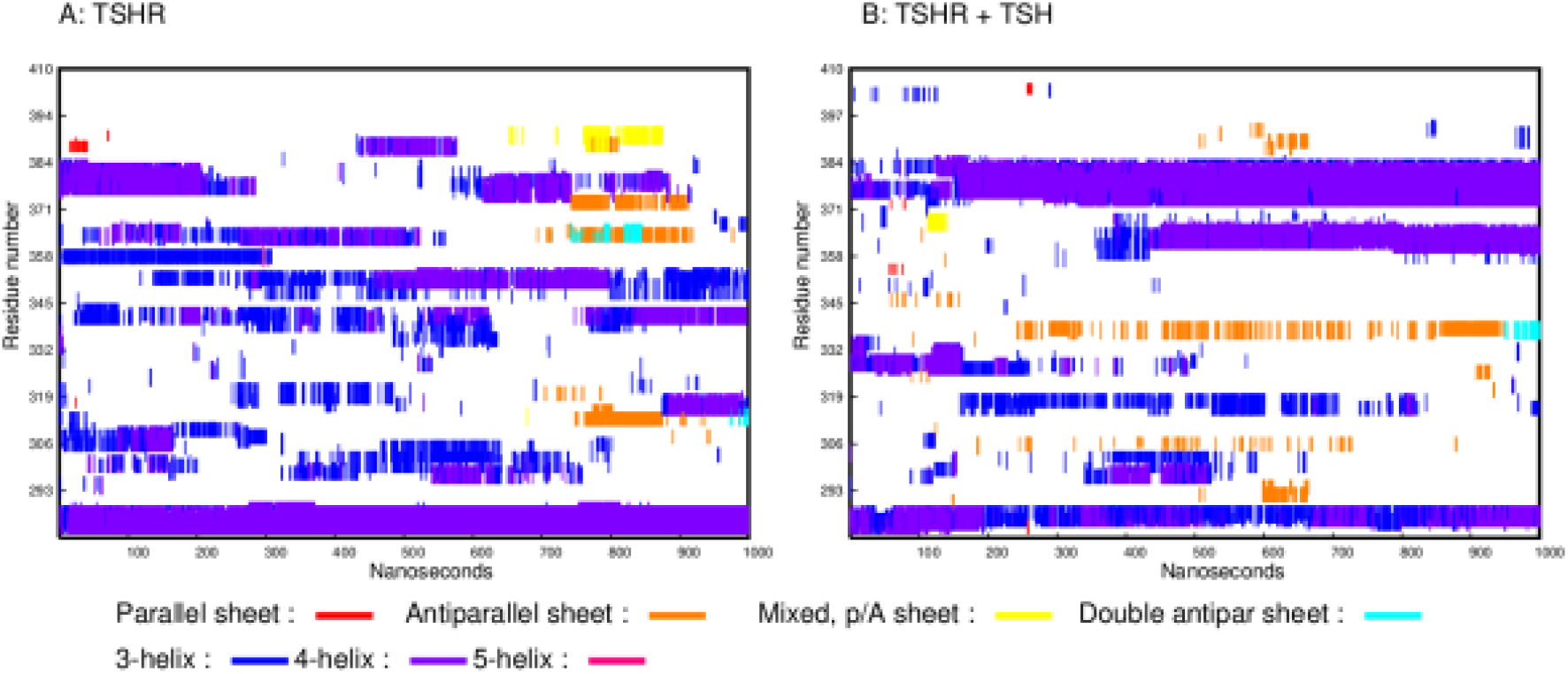
(A) DSSP plot showing the secondary structure elements formed in the LR during the simulation of the TSHR without ligand. The x axis is the simulation time and the Y axis is the residue number. (B) DSSP plot showing the secondary structure elements formed in the LR during the simulation of the TSHR-TSH complex. Compared to (A) there appears to be much improved stabili

The history of hydrogen-bonded residue pairs for the LR is shown in **Figure 7**. Each line on the plot represents one residue pair. By this analysis, it was seen that the inter-domain hydrogen bonds (THR250-VAL374, ALA252-VAL374, LEU254-THR376) persisted throughout the simulation, although several of these residue pairs broke and reformed their hydrogen bonds during the run. This reflected the structural fluctuations similar to the fluctuations seen in the DSSP plot of the SSEs. Note, however, that these hydrogen bonds are not the ones creating most of the SSEs.

**Figure 7:**
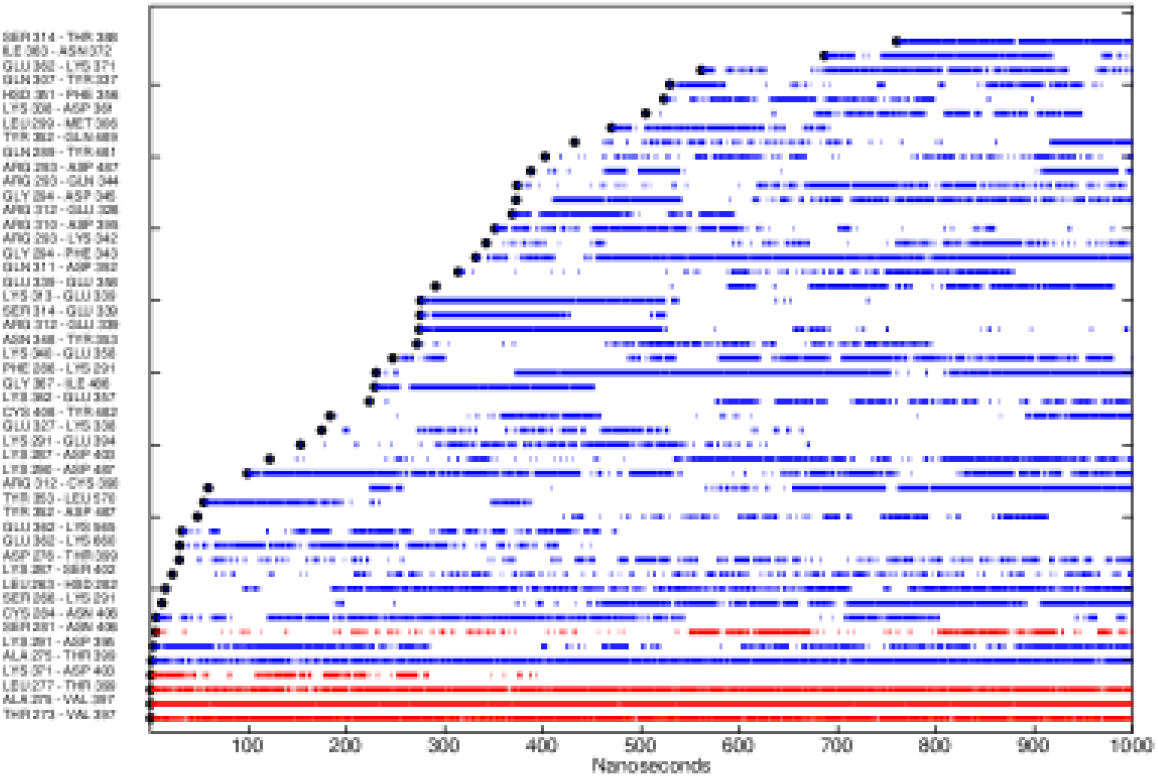
Plot of the residue pairs involving just the LR that were hydrogen bonded at some parts of the simulation. The lines are broken whenever the residue pair was not hydrogen bonded. Blue represents residue pairs within the LR and red represents hydrogen bonds between residues in the LR and the LRD. Note the unbroken lines between the LR and LRD while the LR itself is intrinsically unstable. Note: Residue pairs have to be at least five residues apart (to exclude the many intra-helix hydrogen bonds) and be hydrogen-bonded at least 15% of simulation time to be represented.

### Receptor orientation

During these studies the relative orientation of the LRD with the TMD was also found to undergo significant fluctuations. **Figure 8A** demonstrates this flexibility by showing the conformation of the LRD and LR of the full-length model at 250 ns intervals, superimposed on the initial TMD backbone (without the C-terminal tail). **Figure 8B** shows the fluctuation of the angle between the Z axis and the first principal axis of the LRD. It is also clear from the figures that the rotation of the LRD with respect to the LR is largely confined to one axis. Note also, that all LRD conformations stayed well within the simulation cell. **Figure 8A** also shows the wide range of conformations that the LR forms during the simulation and the changing shape of the 50 AA cleavage region (shown in green) within the LR which is reported to be cleaved by membrane bound matrix metalloprotease (31,32) both in the native and activated states of the receptor (33). Near the end of the simulation, the two end residues of this 50-residue segment (which will form the C peptide) become close (Cα - Cα distance is 6.3 Â) – which may be of significance for post-cleavage processing.

**Figure 8:**
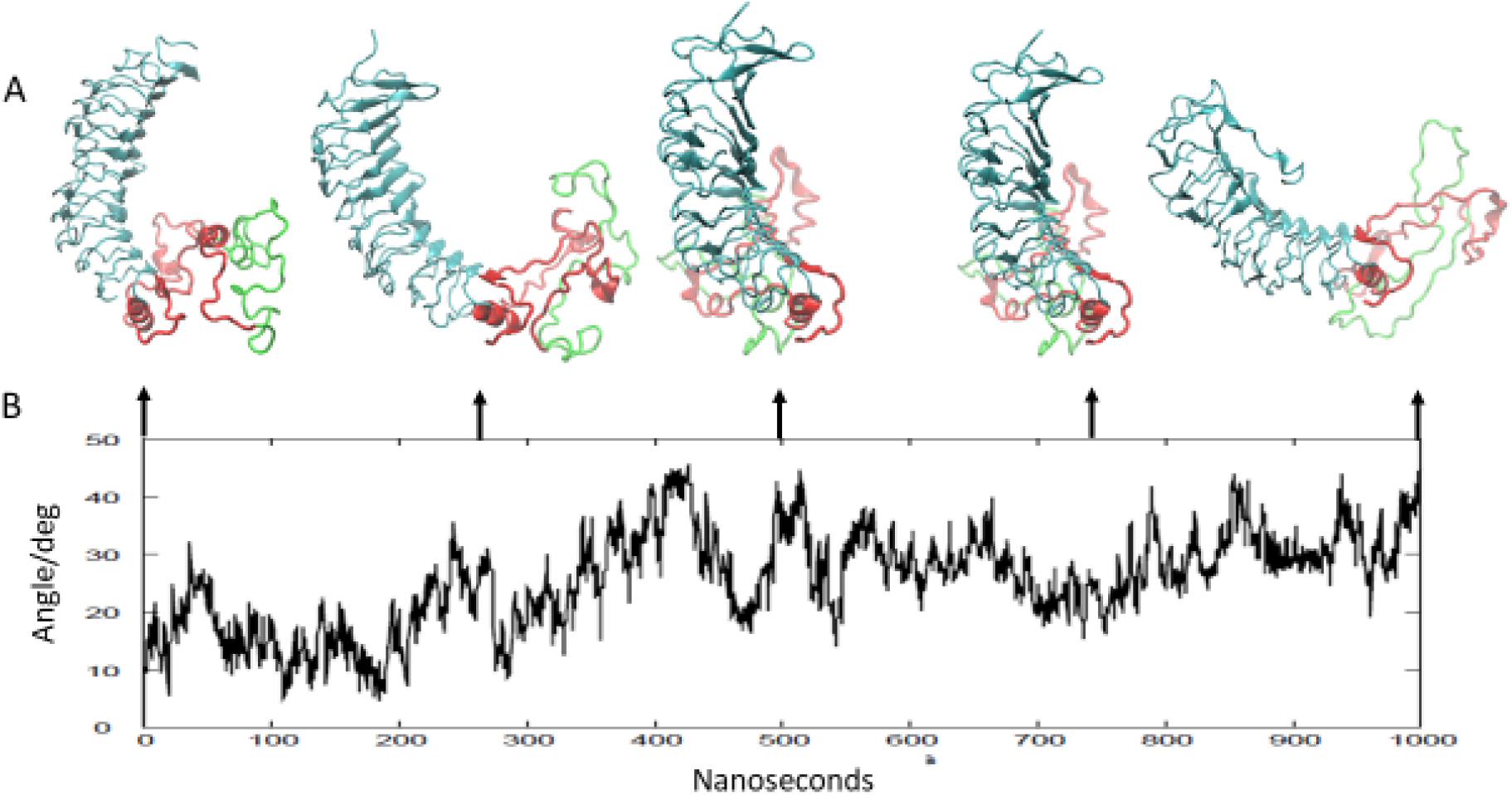
(A) Comparison of the relative orientation of the LRD with the TMD at 250 ns intervals. The structures are aligned by the TMD that is not shown. The LR is shown in red with the 50 AA unique insert (316-366) that may be cleaved during post translational processing is shown in green. (B) The instability of the LR is further illustrated by changes in the angle (in degrees) between the first principal axis of the LRD and the z axis, over 1000 nanoseconds.

### Analysis of the TMD helix bundle

As part of the full-length receptor structure, we also carried out analyses of the constitutive variation in the helix geometry of the 2D RMSD map of 2000 structures. This was calculated based on superimposing the structures on the TMD only as performed earlier. Using k-medoid clustering into three clusters resulted in three representative structures. The changes in the transmembrane helix bundle with the initial model (whose structure was obtained from the third model of the previously published TRIO model (16)) were analyzed with the TRAJELIX (34) module of Simulaid. This program is based on the geometry of the Cα atoms, defining the helix axis (35). Helices with proline are broken up into sub helices; the short segment between the two sub helices in helix 7 is ignored. Data in **Table 1** showed the change in the helix length and in the radius of the circle fitted to the Cα atoms, a measure of the bent of the helix (the smaller it is, the more bent is the helix). The change is the average over the representative structures minus the reference structure’s value. When the reference value is outside the range of the values from the three representative structures, the change is deemed significant. The largest change was observed in **helix 3** that became more curved, resulting in significant shortening (defined as the end-to-end distance).

**Table 1.**
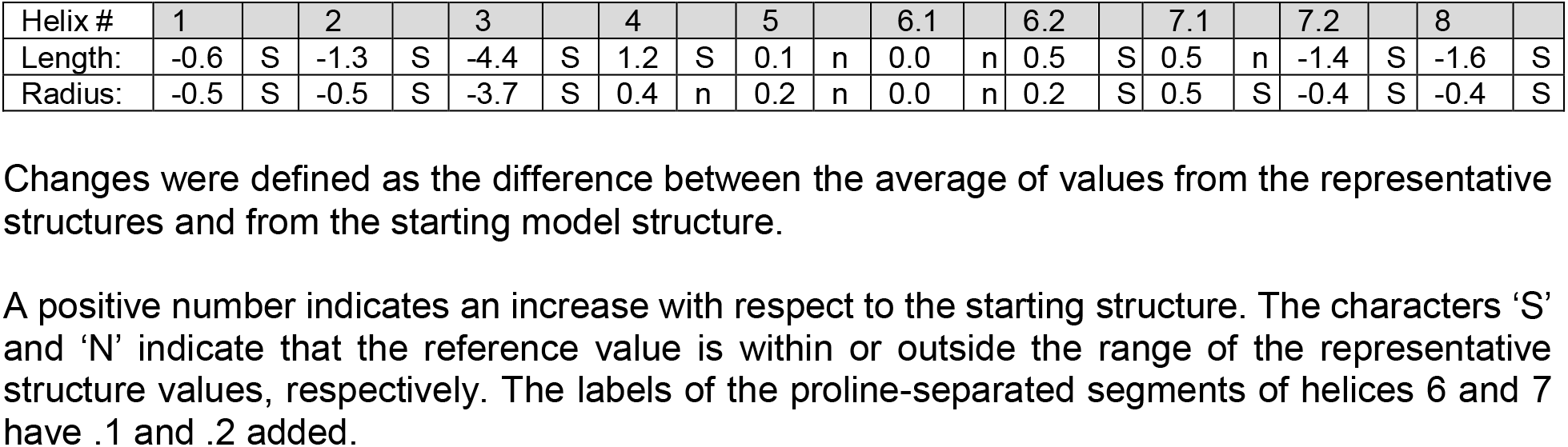
Changes in helix length and radius.

Changes in the distance between the helix centers and the change in the closest approach of the helix axes are shown in **Table 2**. The comparison of the two values gives an indication of the relative shifts. The changes in the helix-helix angles are shown in **Table 3**. We noted from these data that, while the overall arrangement of the helix bundle did not change, it was clear that non-trivial adjustment of the helix bundle occurred. The extent of changes was similar to the changes we observed when the homology model was compared with representative structures from our earlier MD simulation of the TSHR TMD (16).

**Table 2.** Helix-helix distance changes

| Helix# | 1 |  | 2 |  | 3 |  | 4 |  | 5 |  | 6.1 |  | 6.2 |  | 7.1 |  | 7.2 |  | 8 |  |
| --- | --- | --- | --- | --- | --- | --- | --- | --- | --- | --- | --- | --- | --- | --- | --- | --- | --- | --- | --- | --- |
| 1 | 0.0 | n | -0.4 | S | 0.1 | n | -0.1 | n | 2.2 | S | 1.5 | S | 1.9 | n | -5.0 | S | 0.2 | n | -0.4 | n |
| 2 | 0.1 | n | 0.0 | n | 0.4 | S | -1.6 | S | 1.3 | S | -0.9 | S | -0.4 | n | 3.8 | S | 0.8 | n | -5.4 | S |
| 3 | 1.1 | S | 0.7 | S | 0.0 | n | 0.1 | n | -0.4 | n | -1.0 | S | -1.1 | n | 2.3 | n | 7.0 | n | -10.1 | S |
| 4 | 0.5 | n | 0.3 | S | 0.2 | S | 0.0 | n | 0.7 | S | 3.8 | S | -1.3 | S | 8.2 | S | 8.5 | S | -7.2 | S |
| 5 | 2.8 | S | 1.2 | S | -0.2 | n | -0.2 | n | 0.0 | n | -2.0 | S | 0.0 | n | 2.5 | S | 3.2 | n | -11.2 | S |
| 6.1 | 2.7 | S | 0.5 | n | -1.2 | S | -1.1 | S | 0.1 | n | 0.0 | n | -0.8 | S | -1.1 | S | -3.6 | n | -6.9 | S |
| 6.2 | 1.0 | S | -0.8 | S | -1.5 | S | -1.0 | S | 0.1 | n | 0.0 | n | 0.0 | n | -0.3 | n | -2.1 | S | 1.3 | n |
| 7.1 | 2.0 | S | 1.2 | S | 1.2 | S | 1.4 | S | 1.6 | S | 0.7 | S | -0.5 | S | 0.0 | n | 1.0 | n | -7.3 | S |
| 7.2 | 1.9 | S | -2.3 | S | -2.6 | S | -3.7 | S | -0.1 | n | 0.5 | n | 0.2 | n | 1.6 | S | 0.0 | n | -0.9 | S |
| 8 | 1.8 | n | -0.5 | n | -0.8 | S | -2.6 | S | 1.2 | S | 1.9 | S | 0.8 | S | 2.4 | S | 0.3 | n | 0.0 | n |

Upper triangle shows the change in the closest approach of the helix axes; the lower triangle shows the change in the distance between the helix centers. Positive number indicates an increase with respect to the starting structure. The characters 'S' and 'n' indicate that the reference value is within or outside the range of the representative structure values, resp. The labels of the proline-separated segments of helices 6 and 7 have .1 and .2 added.

**Table 3.**
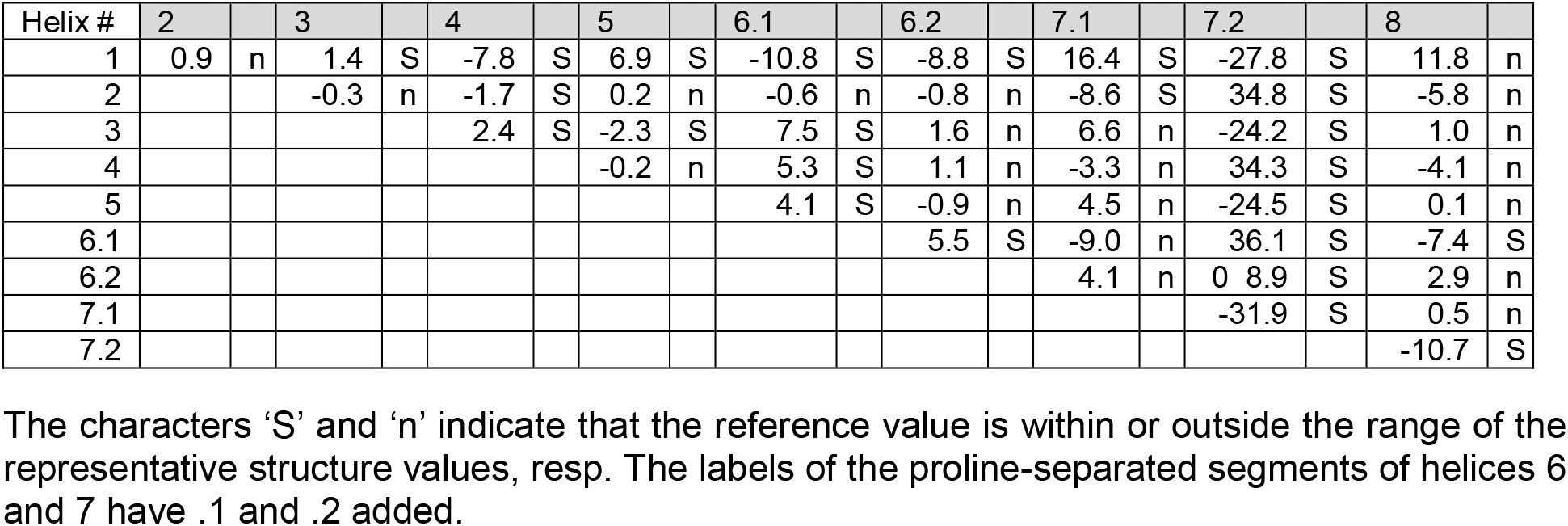
Helix-helix angle changes

### Analysis of the cysteines and cysteine bonds

The LR has six cysteines that are able to form three cysteine bonds: C283-C398, C284-C408, and C301-C390. In fact, in the Alphafold2 structure the distance between the corresponding SG atoms are 3.44, 5.08 and 7.72 Å, respectively. Since the simulation did not include these bonds, it was interesting to see if the LR preferred conformations favorable for the cysteine bonds to form. **Figure 9A** shows the distances for the three putative bonds with time and **Figure 9B** shows the position of the sulfur atoms in these cysteines (colored to match the corresponding graph color) in the Alphafold2 model of the LR. It is clear that C283 and C398 stayed consistently close and C384 and C408 did not separate too far from a bonding distance. However, the third pair, which actually were not close even in the Alphafold2 structure, quickly separated and never became close.

**Figure 9:**
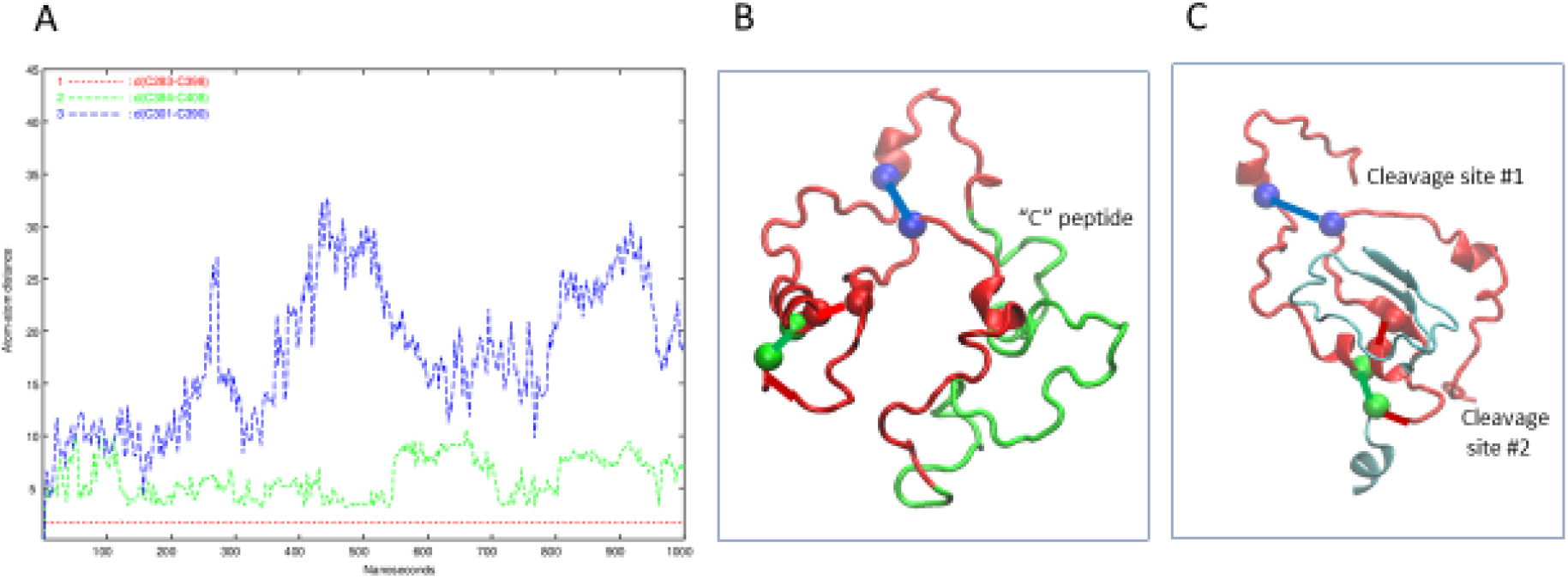
(A) Time evolution of the three cysteine-cysteine distances in the LR. C283-C398: red, C384-C408: green, and C301-C390: blue. (B) The LR backbone (red and green) and the S atoms of the cysteines, colored to match the corresponding graph color. The putative pairs are connected by a line. (C) Here we have left the cysteine pairs connected but taken away the 50 AA insert in the LR and show the reported cleavage sites. For giving context a small part of the LRD and the TMD are also shown in gray/blue.

### Simulation of the TSHR in complex with TSH

Since our conclusion was that the LR is an IDP, it was of interest to find out what, if anything, stabilizes its structure. The most likely candidate was its ligand, TSH. Thus, the structure shown in **Figure 10A** was used to set up an MD simulation modeled after the earlier simulation without the TSH. The DSSP plot of the simulation is shown in **Figure 6B**. The SSEs were remarkably more stable, indicating that the presence of TSH indeed stabilized the LR. The conformations of the complex at the start, middle and end of the 1000 ns simulation are shown in **Figure 10**. It clearly shows that the LR is attached to the TSH at several places. The hydrogen-bond analysis analogous to the one shown in **Figure 7** showed that there are seven residues which are hydrogen bonded to the α subunit of TSH and three LR residues hydrogen bonded to the β subunit and there is even one LR residue that is hydrogen bonded to the LRD. It also shows that 6 of the 10 LR-TSH contacts were formed with the LR residues which are not part of the 50 AA cleaved segment. In addition, new contacts formed between the LR and the LRD with time. On the other hand, the LR moved significantly away from the TMD (more at 500 ns than at 1000ns) indicating that to be able to transmit the signal induced by binding of TSH the cysteine bonds have to be present in the LR. While such analyses remain preliminary and subject to more and possibly longer simulations the structure-inducing effect of the TSH appears to be very clear.

**Figure 10:**
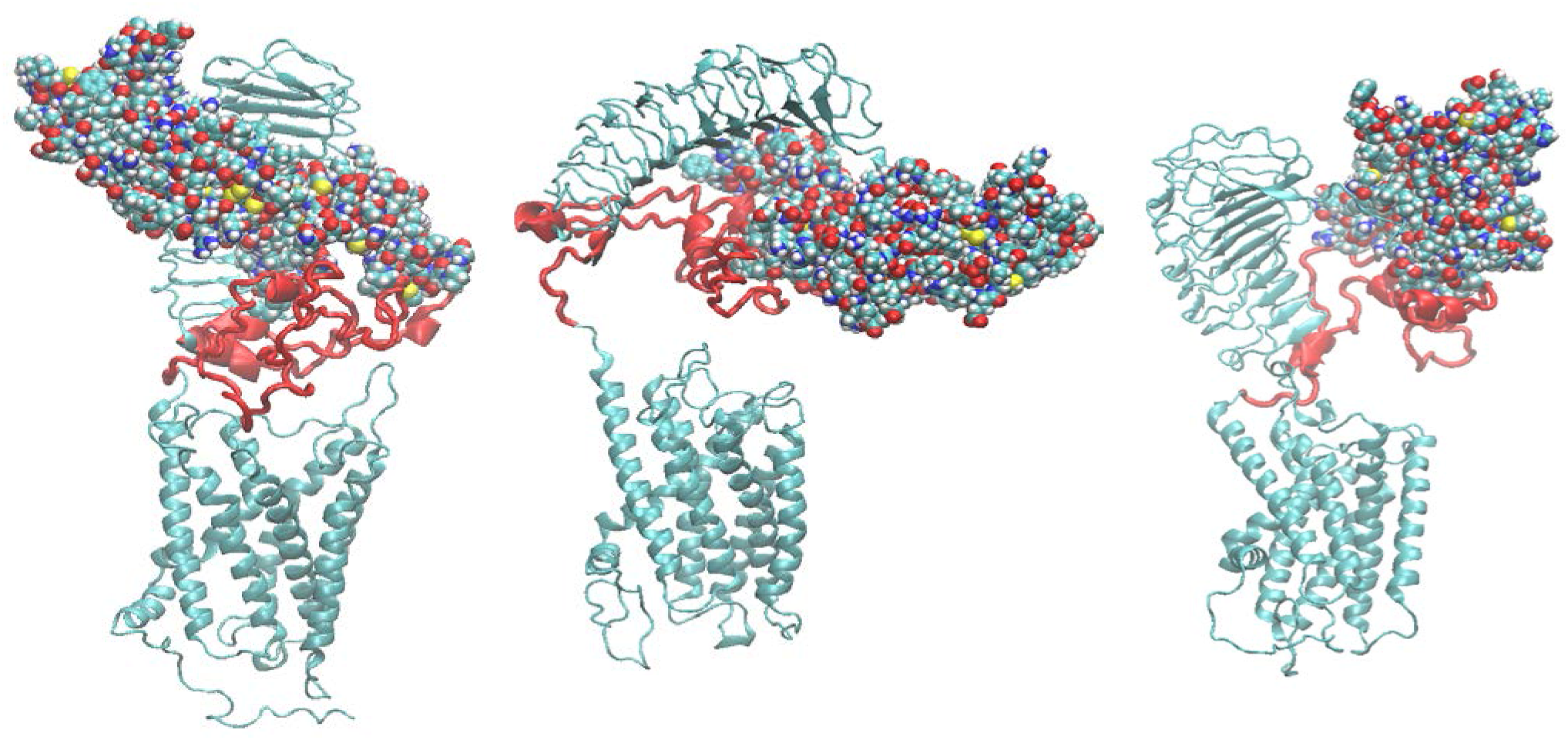
The conformations of the TSHR-TSH complex at the start, middle and end of the simulation. TSH is shown as spheres, the LR backbone is red and the LRR and TMD domains are gray/blue.

## Discussion

The TSHR, similar to the FSH and LH/hCG receptors, consists of a large extracellular ligand bind domain incorporating 11 leucine rich repeats (LRD) and a transmembrane domain (TMD), linked via a 130 AA linker region (LR). The TMD is made up of eight helices joined by extracellular and intracellular loops and a long C-terminal cytoplasmic tail. The TMD is embedded in a phospholipid bilayer and transduces a cascade of signals by engaging several different G proteins (36) and β arrestins (37,38). The interest in the TSHR has been largely fueled by its role as a major human autoantigen in autoimmune thyroid disease; especially Graves’ disease (1).

Detailed mapping of binding sites and interaction partners for the TSHR ligand, TSH, and for stimulating and blocking monoclonal autoantibodies to the TSHR, have been revealed by homology modeling (39) and crystallization of the partial ectodomain bound to these autoantibodies (6,7). Furthermore, homology modeling has suggested possible mechanisms by which activation of the receptor by TSH and a stimulating antibody might occur (40–42). However, all these tripartite models have remained incomplete due to the lack of a reasonable structure for the large TSHR linker/hinge-region (LR). Here we present a full-length model of the TSHR which became possible because of the recent availability of models of the human proteome generated by the artificial intelligence (AI) based protein folding program Alphafold2 (15). We combined the Aplhafold2-generated structure of the LRD – LR complex with our recent MD-refined homology model of the TSHR TMD (16) and further refined the structure with molecular dynamics in a DPCC membrane environment. Given that the structures of the LRD and the TMD of the TSHR have been described in earlier studies (43,44), our analysis in this report was firstly focused on the structure of the LR, its intramolecular and molecular bonding dynamics and its structural variations in lieu with LRD and TMD structures.

It is notable that in the initial structure obtained by the combination of the two parts using the relative orientation of the LRD (residues 22-280) and the TMD (residues 409-694) resulted in a full-length receptor allowing the formation of TMD dimers in a conformation predicted and experimentally verified by our earlier work (44) and which would still leave the LRD binding surface free for ligand or autoantibody binding. However, in the Alphafold2-predicted conformation presented here, the concave surface of the LRD where autoantibodies bind, is partly occluded by the LR and thus would clash with a bound TSHR antibody whose binding sites on the LRD are known (PDB id 3g04). This conundrum was resolved by the observations that (a) the structure of the LR cluster is highly flexible and (b) the relative orientation of the LRD with the rest of the structure is also highly variable resulting in a significant population of conformations where simulating and blocking TSHR autoantibodies could access the receptor to activate or block signaling. Thus, we can say that the Alphafold2 structure of the LR, while useful in providing a starting point for the MD simulation, was not correct in the sense that it missed the large conformational freedom of the LR needed for ligand binding. Given the low reliability score assigned to the LR part of the Alphafold2 structure this observation was not a surprise. However, these conclusions could be further verified by obtaining, if possible, cryo electron microscopy (CEM) or a crystal structure of the native full-length TSHR perhaps stabilized by an autoantibody to the LR.

Based on this study we can conclude that the LR in the TSHR is remarkably flexible, sampling vastly different overall conformations without settling on a well-defined tertiary structure. While different SSEs formed during the simulation (mostly helices), they were transient, as seen from **Figure 6A**. On the other hand, contacts between the LR and the LRD persisted throughout the simulation. The conclusion is that the LR is an IDP. This conclusion is consistent both with the fact that so far no one has succeeded in obtaining its crystal structure and with the suggestion (15) that low reliability scores indicate that the protein is intrinsically disordered. However, from these studies we speculate that the highly flexible nature of the LR is what allows it to accommodate both the ligand or the autoantibodies to the LRD.

The convergence of the simulation is always an open question. However, there are important indicators that suggest that our sampling was adequate. **Figure 7**, the history of hydrogen bonds involving the LR showed that most of the bonds had formed during the first half of the simulation; no new ones formed in the last quarter. Hydrogen bonds also kept forming and reforming, Similarly, **Figure 6A** shows that most SSEs formed and broke several times during the simulation. Taken together these indicators allow us to conclude that the simulation involved adequate sampling.

In addition to our examination of the LR structure, we compared the changes seen in the TMD helix bundle, from the respective reference structure in the TMD-only simulation, versus the full-length model. It was of great interest that the change in the curvature (and, as a consequence, in the end-to-end distance) of Helix 3 was significantly greater in the new full-length model than in the TMD-only model. Comparing the range of values sampled in the representative structures (data not shown) we found similar differences. This observation supported the hypothesis put forward earlier (45) that helix 3 is highly important for the signal transduction of the TSHR and consistent with a variety of small molecule activators which all interact with Helix 3 (not illustrated). We also examined the LR cysteines. Much has been discussed concerning the role of cysteine bonds in anchoring the LR to the LRD following post receptor processing which involves cleavage of the unique 55 amino acid insert in the LR (see **Figure 9C**). The analysis of the cysteines in the LR showed the remarkable affinity of one pair to each other (C283-C398) and to a lesser degree for the second pair. (C384-C408). However, the third pair (C301-C390), showed poor affinity, leading to the suggestion that in the fully formed TSHR only two of the cysteine bonds were formed.

Recently, structures of the luteinizing hormone receptor (LHR) in complex with different proteins and in its inactive state, a total of five structures, obtained by CEM have been described (46). In accordance with our simulation, the relative orientation of the LRD with the TMD was quite variable. The LR in the LHR, however, while significantly shorter than the LR in the TSHR, was missing more than 40 residues in the CEM structures thus preventing detailed comparison with the TSHR LR. It is notable that the short helix at the N-terminal end of the LR was present in all five CEM structures and during the 1000 ns MD run described here. Also, the LR conformations in the CEM structures were very different from each other, reflecting the conformational variations observed during our TSHR MD run. Interestingly, our simulation of 1000ns with the heterodimeric TSH showed stabilization of secondary structural elements which was clearly absent without TSH (**Figure 6**) further suggesting that the LR plays an important part in TSH binding and that TSH is able to stabilize the flexible LR region after transitioning through various dynamic structural changes. Furthermore, the separation of the LR and the TMD during the simulation suggests that for the proper action of the TSHR on binding of the TSH then at least one cysteine bond has to be formed.

In conclusion, we generated the first model of the full-length TSHR that includes the characterization of a flexible LR and concluded that the LR is constitutively unstable in the native state of receptor thus can be considered an IDP. However, our preliminary results indicate that TSH is able to stabilize this disordered protein, which suggest that stabilization of LR might be important for signaling to ensue.

## Additional information

### Disclosures

TFD is member of the Board of Kronus Inc (Starr, ID, USA); MM, and RF have nothing to disclose.

### Data Availability

The initial model generated is available from the Dryad server; URL: https://doi.org/10.5061/dryad.rjdfn2zdp.

## Abbreviations

AA: amino acid
CEM: cryo electron microscopy
DPPC: Dipalmitoylphosphatidylcholine
ECD: ectodomain
GCE: grand-canonical ensemble
GPCR: G protein coupled receptor
IDP: intrinsically disordered protein
LHR: luteinizing hormone receptor
LR: linker region
LRD: leucine-rich domain
MD: Molecular Dynamics
SSE: secondary structure element
TMD: trans-membrane domain
TSH: thyroid stimulating hormone
TSHR: TSH receptor

## Acknowledgments

This work was supported in part by NIH grant DK069713 and a VA Merit Award BX000800 (to TFD), the Segal Family Fund and additional anonymous donations. It was also supported in part through the computational resources and staff expertise provided by the Department of Scientific Computing at the Icahn School of Medicine at Mount Sinai.

---

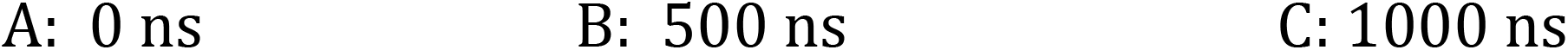

